# Co-evolutionary Distance Prediction for Flexibility Prediction

**DOI:** 10.1101/2020.10.15.340752

**Authors:** Dominik Schwarz, Guy Georges, Sebastian Kelm, Jiye Shi, Anna Vangone, Charlotte M. Deane

## Abstract

Co-evolution analysis can be used to accurately predict residue-residue contacts from multiple sequence alignments. The introduction of machine-learning techniques has enabled substantial improvements in precision and a shift from predicting binary contacts to predicting distances between pairs of residues. These developments have significantly improved the accuracy of *de novo* prediction of static protein structures. Here we examine the potential of these residue-residue distance predictions to predict protein flexibility rather than static structure. We used DMPfold to predict distance distributions for every residue pair in a set of proteins that showed both rigid and flexible behaviour. Residue pairs that were in contact in at least one reference structure were considered and classified as rigid, flexible or neither. The predicted distance distribution of each residue pair was analysed for local maxima of probability indicating the most likely distance or distances between a pair of residues. The average number of local maxima per residue pair was found to be different between the sets of rigid and flexible residue pairs. Flexible residue pairs more often had multiple local maxima in their predicted distance distribution than rigid residue pairs suggesting that the shape of predicted distance distributions is predictive of rigidity or flexibility of residue pairs.

## Introduction

Important functions of proteins like catalysis or molecular recognition are often coupled to switching between multiple structurally distinct states of protein complexes or single proteins.^1, 2^ Here we examine the structural changes of single polypeptide chains. Such changes can range from side chain movements to whole domain movements^1^ and are always associated with specific residues or regions being flexible. Knowledge about residue flexibility or about a protein’s structural ensemble would improve our mechanistic understanding of protein function and benefit areas such as *in silico* screening for drug discovery.^3^

Protein flexibility can be experimentally determined through techniques such as Nuclear magnetic resonance (NMR)^4^, Förster resonance energy transfer (FRET)^5^, Hydrogen deuterium exchange (HDX)^6, 7^ or X-ray free-electron laser (XFEL)^8^ experiments. While NMR spectroscopy has been widely used to gain structural and dynamic information about proteins, the technical difficulties increase for proteins larger than 20kDa (about 200 amino acids).^4^ Although protein size is less of an issue for FRET, HDX and XFEL studies, all these techniques require extensive experimental resources making large-scale application infeasible.

Due to these limitations, several methods have been developed for the computational prediction of protein flexibility. They can be divided into two main types: prediction of residue flexibility and prediction of a protein’s structural ensemble. Individual residue flexibility can be predicted using many techniques including information from crystallographic B-factors^9^, normal mode analysis (NMA)^10^, NMR chemical shift data^11^ and Molecular Dynamics (MD) simulation data^12^. While some of these methods can predict residue flexibility from sequence alone, the predictions contain no information about different distinct conformations of a protein. In contrast, ensemble prediction methods such as NMA^13, 14^, distance geometry^15^ or coarse-grained MD simulations^16^ generate information about distinct conformations of a protein but currently no ensemble predictor uses only sequences as input. In fact, most of the methods that predict individual residue flexibility and all of the ensemble predictors rely on experimentally determined or fully modelled structures to guide analysis. Since sequence information is generally more widely available than accurate structural data, it would be desirable to have ensemble predictors that only take protein sequences as input. This would enable the large-scale prediction of structural ensembles at low computational cost.

Directly predicting multiple structurally distinct states of a protein is far more challenging than predicting individual residue flexibility but it offers more insights and direct applicability in further research studies such as *in silico* screening. We introduce a third concept of protein flexibility that might bridge this gap between residue flexibility and ensembles: residue pair flexibility. It describes the flexibility of residue-residue interactions that change when proteins switch between different structurally distinct states to perform their function. Knowledge about the rigidity or flexibility of two residues’ interactions can inform the prediction of a protein’s structural ensemble. Prediction of residue-residue interaction, in particular the prediction of residue-residue distances, has recently been developed and applied in static protein structure prediction.^17–20^ Co-evolutionary distance prediction is a method that uses only sequences as input. We examined whether this method could also predict residue pair flexibility and thus, could potentially be applied to ensemble prediction.

Co-evolutionary contact predictions have been used to improve *ab initio* prediction of static globular protein structures^21^, domain boundaries^22^, protein-protein interactions^23^, and loop structures^24^. Additional improvements have been made by training neural networks on large sets of available protein structures alongside co-evolutionary information.^25–27^ Most recently these data have been used to train methods to predict residue-residue distances instead of binary contacts.^17–20^ These distance predictions yield a probability distribution over distance bins, with local maxima of probability indicating the most likely distance or distances between pairs of residues. For the earlier methods that predicted only binary contacts^28–30^, it was found that when two residues were in contact across all known PDB^31^ structures of a protein sequence, the predicted residue-residue coupling score was ranked higher than a residue pair that was only in contact in a subset of the structures of that protein sequence.^32^ Since in these binary methods a lower rank indicates a lower probability, residue-residue contacts that only exist in some states of a protein’s ensemble were more likely to be falsely classified as not in contact (at all). We wanted to test whether the recent improvements in co-evolution analysis and distance predictions encode information about different conformers of a protein in the shape of their predicted distance distributions. We hypothesise that predicted distance distributions with more than one local maximum indicate residue pairs that can adopt more than one metastable state, and that having more than one local maximum is therefore related to flexibility.

As mentioned above, flexibility can be determined by different experimental or computational methods which each have their limitations, especially for large sets of proteins. Another way of approximating the flexible nature of a protein is the comparison of its different available PDB structures.^33, 34^ These static representations of proteins lack the dynamic component of flexibility but represent multiple structurally distinct conformations of a protein’s ensemble and thus, approximate the degree of flexibility and the change of interactions between pairs of residues. Here, we use the database of Conformational Diversity in the Native State of proteins (CoDNaS), which stores multiple PDB structures of a single protein sequence.^34^ For our analysis we used the maximum RMSD pairs dataset of Monzon *et al.*^35^, a subset of CoDNaS containing two PDB structures of one sequence that showed the maximum RMSD between all pairwise comparisons of a protein’s available PDB structures. These structure pairs are a proxy for a protein’s ensemble of structurally distinct states and can be used to estimate and classify residue pair flexibility.

We find that the number of local maxima in a residue pair’s predicted distance distribution is different between residue pairs that are rigid or flexible. We calculated the number of local maxima in the predicted distance distribution of each residue pair indicating if a residue pair has a single or multiple probable distances predicted. We found that flexible residue pairs had a significantly higher fraction of multiple local maxima than rigid residue pairs, implying that flexible residue pairs were predicted to be in more than one metastable state more often than rigid residue pairs. We show that this difference was not biased by individual proteins and that it was also not driven by the different secondary structure proportions which were present between the sets of rigid and flexible residue pairs. Furthermore, we observe very similar local maxima counts to the CoDNaS dataset in the predicted distance distributions of a second dataset containing a set of rigid loops validated by MD simulations. We therefore concluded that the shape of predicted distance distributions is informative of stability or flexibility of residue pairs and could potentially be used in protein ensemble prediction.

## Results & Discussion

### The shape of predicted distance distributions is related to flexibility

Co-evolutionary information and machine-learning have recently enabled the prediction of residue-residue distances.^17–20^ These methods yield a predicted distance distribution for every residue pair and have been used to identify the most likely distance (or distance interval) between a pair of residues in a protein. Such distance constraints have been successfully used to predict static protein structures, e.g. Kryshtafovych *et al.*^36^.

Here we test whether these predicted distance distributions also contain information about residue pair flexibility. Information on which residues change their interactions to facilitate the switching between conformers will be crucial to predict multiple biologically relevant conformations. To identify these interaction changes we analysed and compared the two most different PDB structures of a protein sequence. All residue pairs that are present in both of these structures were classified into rigid, flexible or neither (only non-trivial, true contacts considered; See Methods for more information).

All analysed residue pairs are present in two conformations (two distinct PDB structures), so that for each residue pair two C_*β*_-C_*β*_ distances and thus, a C_*β*_-C_*β*_ distance difference could be determined (C_*α*_ for glycine). Rigid residue pairs were defined as those with a C_*β*_-C_*β*_ distance difference below 1Å and flexible pairs were those with a C_*β*_-C_*β*_ distance difference equal or above 2Å. Residue pairs with a distance difference between these two cutoffs were not classified into either set. Additional to the distance difference thresholds we applied a bond change criterion. Rigid residue pairs had to have at least one physicochemical bond in both structures and flexible pairs had to display at least one bond in one structure but none in the other. This criterion was applied to ensure interaction change for residue pairs that were classified as flexible. More details on the rigid and flexible definitions can be found in Methods.

If the shape of the predicted distance distributions relates to the conformational ensemble of a protein, rigid residue pairs should be characterised by predicted distance distributions with only one local maximum and flexible pairs should show predictions with two or more local maxima.

Figure 1 illustrates the expected difference between rigid and flexible residue pairs. The two residues of the rigid residue pair remain in the same orientation to one another in the two structures (Figure 1a), the two residues of the flexible pair have changed orientations between the two structures (Figure 1b). The distance prediction of the rigid pair shows a distribution with only one local maximum and the two C_*β*_-C_*β*_ distances of the two structures are close to that maximum (Figure 1c). In contrast to that, the flexible pair’s distribution shows two local maxima and the C_*β*_-C_*β*_ distances of the two structures differ by more than 4Å (Figure 1d).

**Figure 1.**
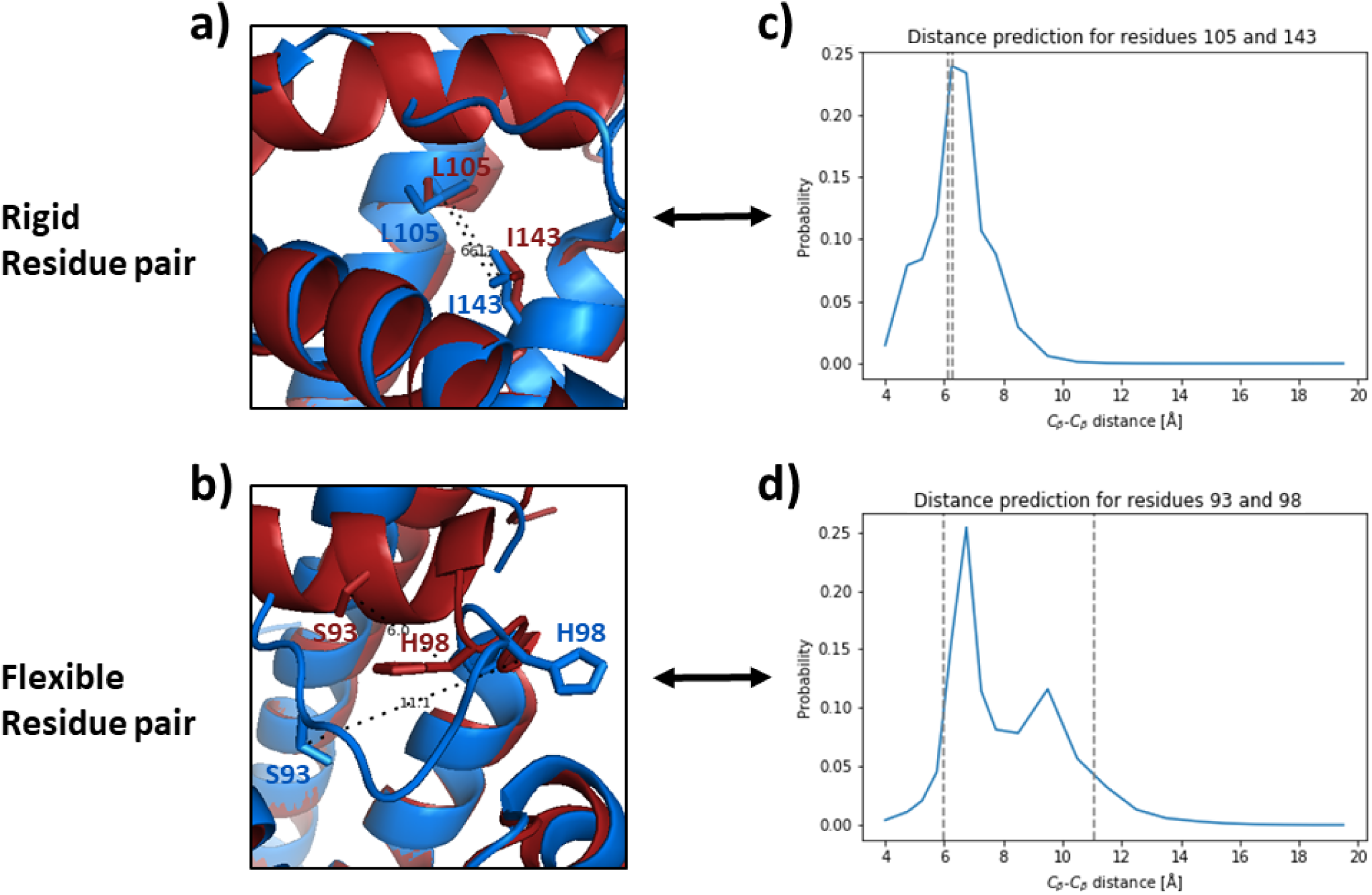
A rigid residue pair associated with a single local maximum and a flexible residue pair with multiple local maxima in their predicted distance distributions are shown. **a)** displays an example of a rigid residue pair of Myoglobin (L105:I143) and **b)** a flexible pair (S93:H98). The two most different PDB structures of Myoglobin were superimposed and in both images PDB 2EB8 A is shown in blue, 2JHO A in red and the residue pairs in stick representation. **c)** and **d)** depict the predicted distance distributions for the respective residue pairs. Black dotted lines in PDB structures as well as vertical dotted lines in probability distributions represent the two C_*β*_-C_*β*_ distances that are associated with the residue pairs in both PDB structures. The rigid residue pair in **a)** is very similar in both structures and hence, its two C_*β*_-C_*β*_ distances are close to each other in **c)** next to the single local maximum of the distance prediction. The flexible residue pair in **b)** displays much greater movement and two C_*β*_-C_*β*_ distances greater than 4Å apart which roughly represents the difference between the distance prediction’s two local maxima in **d)**.

### Flexible residue pairs have more distance prediction local maxima than rigid residue pairs

To test whether there is a general relationship between the flexibility of a residue pair and its predicted distance distribution, we investigated 2947 proteins from the CoDNaS database. We classified each residue pair in each of those proteins into rigid or flexible (or none) and examined the predicted distance distributions for all residue pairs in those sets. Local maxima in the predicted distance distributions were defined as bins that were followed by a local minimum of at least 3% probability below its maximum value (see Methods). For a given set of residue pairs the number of local maxima in each predicted distance distribution was determined and from all those predictions the fraction of predictions with no, one, two or more local maxima was calculated. The different fractions of local maxima counts for all residue pairs, rigid residue pairs and flexible residue pairs across all proteins are shown in Figure 2a.

**Figure 2.**
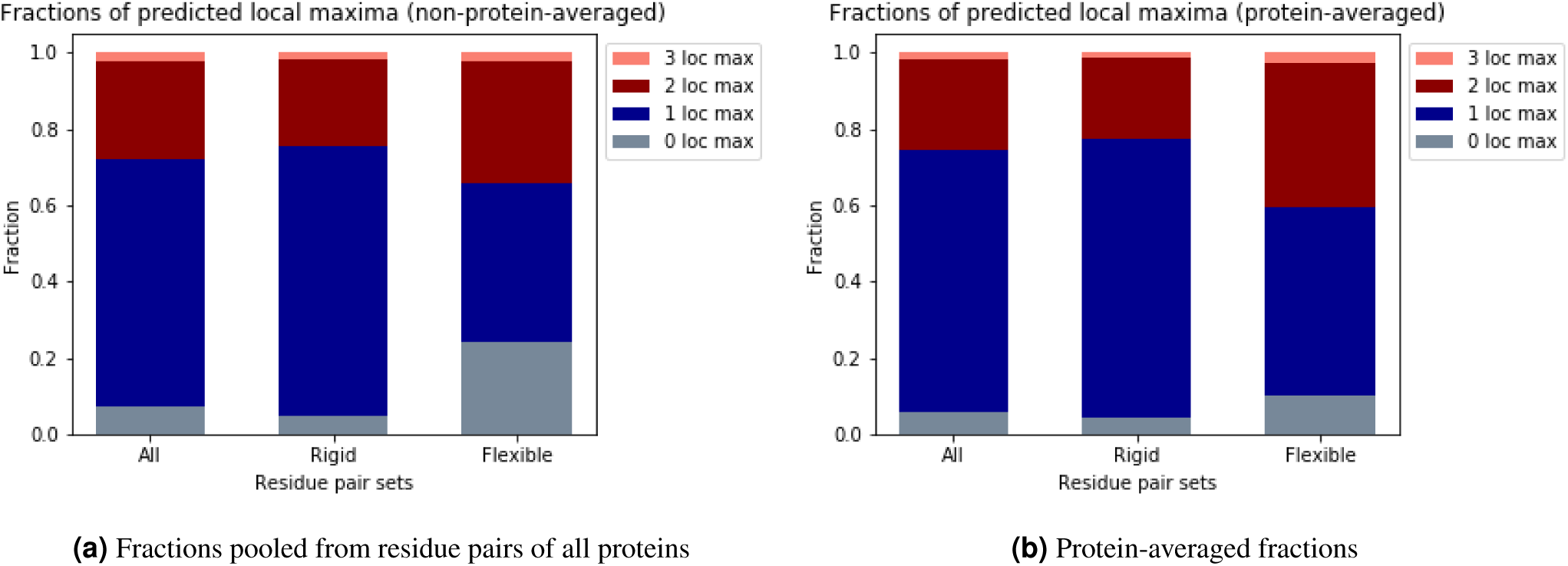
Fractions of local maxima counts are different between rigid and flexible residue pairs. **(a)** Fractions of predicted local maxima across all classified residue pairs of all proteins, showing that flexible residue pairs are both more likely to have no local maximum or multiple local maxima. **(b)** Protein-averaged fractions of predicted local maxima, showing the same overrepresentation of no or multiple local maxima for flexible residue pairs.

Cases where no local maximum could be detected in a distance prediction (grey) implied that either both residues were predicted to be further than 19Å apart or that the predictor did not predict a highly probable distance (see Methods for definitions), 7% of all residue pairs that we investigated fell into this category. In the set of rigid residue pairs only 5% showed no local maximum but in the set of flexible residue pairs 24%, indicating that predicting residue-residue distances for flexible residue pairs is more challenging. 70% of rigid residue pairs had a single local maximum, whereas this was only 42% for flexible residue pairs.

As there are different numbers of residue pairs in the respective sets: 1,370,737 pairs in total, 711,239 in the rigid set and 8,681 in the flexible set and to show that no individual proteins are driving the differences in the number of local maxima, protein-averaging was performed. For each protein the number of local maxima was calculated for each residue pair (’all’) and for the sets of rigid and flexible residue pairs if a protein had at least one residue pair with a single local maximum and at least one other pair with multiple local maxima. Protein-averaged fractions for the residue pair sets are shown in Figure 2b. On average, of all residue pairs 25% ± 8% had multiple local maxima predicted. A fraction of 22% ± 7% of the rigid pairs were found to have multiple local maxima compared to 40% ± 17% of the flexible ones. The difference in the fraction of multiple local maxima between rigid and flexible residue pairs is slightly higher when applying protein-averaging but qualitatively the results are the same (Figures 2a and 2b).

This is also true when computing the local maxima fractions for a subset of proteins that consist only of unique CATH superfamilies (see SI Figure 2). The differences between rigid and flexible residue pairs are qualitatively the same as well.

### Predicted distance distributions can capture flexibility independent of secondary structure

Table 1 shows that the residues in pairs that were classified as flexible were more often part of loop structures than the residues in rigid residue pairs (85% vs. 42%). This is expected since loop structures are often intrinsically flexible^37^. As it is known that loop structures are hard to predict^24^, a higher fraction of multiple local maxima could just stem from an increased uncertainty of predictions and not from residue pair flexibility that relates to distinct conformations. This would imply that the higher fraction of multiple local maxima observed in flexible residue pairs compared to rigid pairs is due to imbalances in secondary structure type proportions and not driven by residue pair flexibility. Therefore, we analysed the secondary structure composition of all sets of residue pairs and the local maxima fractions of different secondary structure subgroups within the two sets of interest. If the local maxima fractions in the respective rigid and flexible secondary structure subgroups were identical and the overall differences were only caused by different proportions of subgroups, the number of local maxima would predict secondary structure type and not flexibility itself.

**Table 1.** Fraction of secondary structure classification depending on flexibility classification

|  | Within/between secondary structure elements | Including at least one loop/coil residue | Proteins with at least one classified pair |
| --- | --- | --- | --- |
| All residue pairs | $0.52 \pm 0.13$ | $0.48 \pm 0.13$ | 2934 |
| Rigid pairs | $0.58 \pm 0.12$ | $0.42 \pm 0.12$ | 2914 |
| Flexible pairs | $0.15 \pm 0.14$ | $0.85 \pm 0.27$ | 1584 |

Figure 3 shows the local maxima fractions dependent on secondary structure subgroups comparing the sets of rigid and flexible residue pairs. Even when stratifying by secondary structure, we still observe differences between the number of local maxima of the rigid and flexible sets. Whereas the subgroups with at least one residue being part of a loop (S-L and L-L) of rigid and flexible sets show similar fractions of multiple local maxima (Figures 3b and 3c), the subgroups where both residues of a pair are part of a secondary structure element (S-S) differ with flexible residue pairs having about twice as many predictions with multiple local maxima compared to the rigid pairs (red in Figure 3a). This, and the large differences in single local maximum fractions (blue) in all subgroups, confirms that the differences between rigid and flexible residue pairs are not driven only by different proportions of secondary structure in the sets.

**Figure 3.**
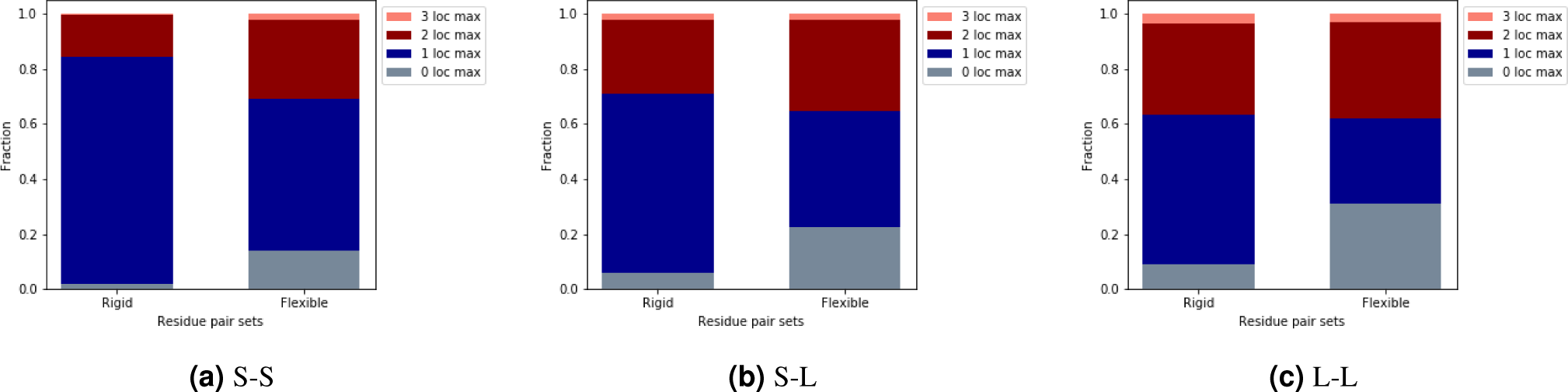
Predicted distance distributions capture flexibility independent of secondary structure. Number of local maxima fractions were computed for subgroups depending on their secondary structure annotation: ‘S-S’ when both residues were found to be part of a secondary structure element, ‘S-L’ when one of the residues was in a secondary structure element and the other in a loop, and ‘L-L’ when both were part of a loop/coil region. The different fractions of multiple local maxima (red) between rigid and flexible residue pairs are not only driven by an imbalance of secondary structure element and loop residues: for example the S-S subgroup of the flexible set has a fraction of multiple local maxima twice as big as the rigid set’s.

As for the overall fractions, we also computed local maxima fractions of the secondary structure subgroups for a subset of proteins with unique CATH superfamilies (see SI Figure 3). The differences in secondary structure subgroups between rigid and flexible residue pairs are also found when controlling for a potential superfamily bias which further supported our finding. XXchange SI figure numbers in manuscriptXX

Our definition of rigid depends on the currently solved crystal structures, so it is possible that when a residue pair is classified as rigid it might not be, alternative conformations might not yet have been solved. However, if a residue pair is classified as flexible, this is likely to be correct as it has already been found in at least two different distinct conformations. As residue pairs are more prone to be incorrectly classified as rigid than as flexible, the fraction of multiple local maxima of rigid residue pairs is likely to be rather overestimated than underestimated. This implies that the difference in multiple local maxima fractions between rigid and flexible residue pairs could increase with more structural data becoming available, further supporting our finding that residue pair flexibility is captured independent of secondary structure imbalances.

### Differences in number of local maxima between rigid and flexible sets are statistically significant

Given that these differences in predicted distance distributions are related to flexibility and rigidity of residue pairs, we tested if these differences are statistically significant. For this we performed protein averaging and analysed the fraction-per-protein distributions. For each protein the fraction of all residue pairs with multiple local maxima was determined and as well as the fraction within the sets of flexible and rigid residue pairs, respectively (See SI for distribution histograms).

The number of proteins contributing to the distribution of all proteins was 2899, 2858 for the rigid sets and 542 for the flexible sets. Note the differences to the numbers when comparing secondary structure pairs in Table 1 (1584 vs. 542). Applying a criterion of having at least one residue pair with a single local maximum and at least one other pair with multiple local maxima in a given protein reduced the number of proteins by about two thirds. We applied this criterion to reduce the potential bias from proteins with a very low number of classified residue pairs.

All three distributions are significantly different from one another (Mann-Whitney U test): all vs. rigid sets with a p-value *<* 10^-44^, all vs. flexible sets with a p-value *<* 10^-87^ and rigid vs. flexible with a p-value *<* 10^-122^. These results not only show that the shape of predicted distance distributions of rigid and flexible residue pairs differ significantly, but also that, when comparing to rigid residue pairs, flexible residue pairs more often have multiple (instead of single) local maxima in their predicted distance distributions.

### Examining the local maxima fractions on a set of protein loops defined as rigid

In order to see if our findings generalised to other methods of identifying protein flexibility, we investigated a set of 20 loops that had been found to move very little during MD simulations^38^. In a large-scale flexibility analysis of MD simulations of 250 proteins these loops had a mean movement of less than 0.5Å and thus, were at the very rigid end of the spectrum of all loops analysed.

We computed the number of predicted local maxima for each residue pair that involved at least one of the loop residues of these 20 rigid loops (496 pairs in total) and generated per-rigid-loop-averaged fractions. This rigid loop set of residue pairs had on average 72% ± 23% of predicted distance distributions with a single local maximum, 25% ± 13% with multiple maxima and 3% ± 10% without any local maximum. Since the loops of this set had shown little flexibility during simulations, their fractions should be similar to the rigid residue pairs of the CoDNaS set which is indeed what we observe (see Figures 4a and 4b for a comparision of S-L and L-L secondary structure subgroups). Although generally similar, the fractions of multiple local maxima are even lower and the fractions of single local maxima are even higher in the rigid loops than in the rigid residue pairs of the CoDNaS set. These results agree with the idea described above that some rigid residue pairs within the CoDNaS set might be incorrectly classified as rigid because alternative conformations have not yet been observed in crystal structures.

**Figure 4.**
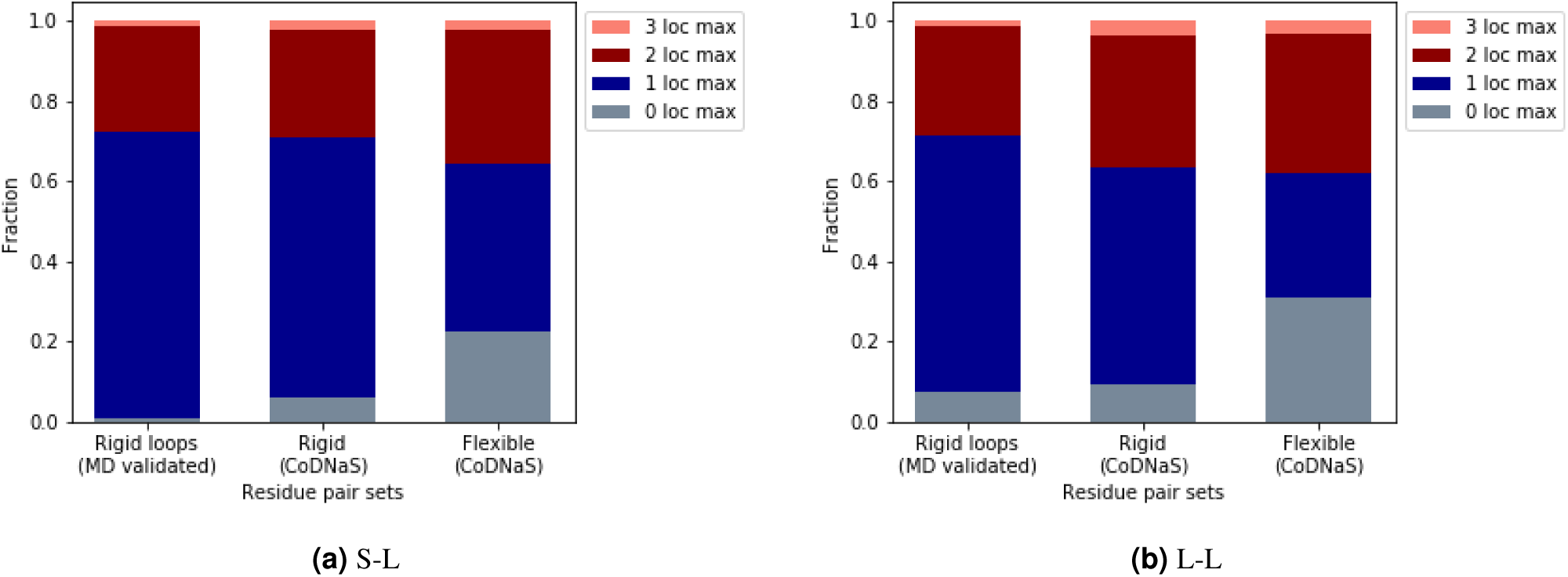
Local maxima fractions of the rigid loop set are similar to fractions of the rigid residue pairs of the CoDNaS set. Secondary structure subgroups ‘S-L’ and ‘L-L’ show small fractions of distance predictions with multiple local maxima (red) and large fractions with a single local maximum (blue). The fractions of multiple local maxima are generally similar to the fractions of rigid residue pairs in the CoDNaS set. This rigid loop set that was defined not by comparing two PDB structures but by analysing MD simulations, confirms the finding that rigid residue pairs less often have multiple local maxima in their distance predictions than flexible pairs.

This set of rigid loops was defined by an alternative validation strategy, not by comparing two PDB structures but by analysing MD simulations, and shows once again that rigid residue pairs have multiple local maxima predicted less often than flexible pairs.

## Conclusion

Switching between different structurally distinct states is common in proteins carrying out various functions like catalysis or molecular recognition. For this switching between conformations, flexibility and the change of specific residue interactions is necessary. Since non-machine learning contact predictors tend to simply assign lower probabilities to such interaction-changing residue pairs (see SI and Zea *et al.*^32^), we wanted to test if recently developed methods which predict residue-residue distances capture information about these changing interactions in a more exploitable way. Specifically, in this study we investigated if rigid and flexible residue pairs have different numbers of local maxima in their predicted distance distributions. For this we analysed 2947 proteins of the CoDNaS database. For every protein the two most different available PDB structures were taken and residue pairs that were present in both structures were grouped into rigid and flexible sets. We analysed and defined residue pair flexibility according to the C_*β*_-C_*β*_ distance difference between the two different conformations of a residue pair and added a bond change criterion to ensure interaction change.

When analysing the number of local maxima in the predicted distance distributions of rigid and flexible residue pairs, the two groups were found to have different fractions of single and multiple local maxima. Rigid residue pairs displayed a larger fraction of single local maxima than flexible pairs whereas flexible residue pairs had a larger fraction of multiple local maxima than rigid residue pairs. As each local maximum is potentially related to a distinct distance that stems from a biologically relevant conformation of a protein (see Figure 1), these results imply that distance predictions capture information about residue pair flexibility and therefore different protein conformations.

Since flexible residue pairs are more commonly found as part of loop structures which are often intrinsically flexible, we tested if this imbalance of secondary structure proportions between rigid and flexible sets was the sole driver of differences between residue pair sets. We found that this was not the case and the difference between sets was also present in secondary structure subgroups. Hence, the information on residue pair flexibility that is captured by distance predictions is not only due to the uncertainty of predictions for areas of loop/coil secondary structure.

After confirming that distance predictions capture residue pair flexibility and not only secondary structure, we performed protein averaging of the fractions of single and multiple local maxima. This yielded distributions of fractions on a per-protein basis which allowed us to demonstrate that the differences between rigid and flexible residue pairs are statistically significant.

We then tested if our results held when considering protein flexibility as defined by Molecular Dynamics. We predicted distance distributions for all residue pairs within and in contact with a set of loops which were found to be rigid in MD simulations. Although the validation of rigidity differed from our approach of comparing two PDB structures of one protein, the resulting fractions of single and multiple local maxima were similar to the set of rigid residue pairs in the CoDNaS set. This alternative dataset supports our findings of differences in distance predictions of rigid and flexible residue pairs. As discussed above, some rigid residue pairs might have been incorrectly classified as rigid because alternative protein conformations, that might show a flexible behaviour of that residue pair, have not yet been solved. The even higher fraction of predicted distance distributions with a single local maximum of residue pairs that were classified as rigid by MD simulations compared to residue pairs classified by PDB structure pairs further strengthens this view.

The fact that the average number of local maxima and thus, the number of predicted distances that are associated with these local maxima varies between rigid and flexible residue pairs could be used to improve multi-conformer modelling. Current structure prediction tools aim to predict the static structure of a protein^17–20^ and multiple output structures are an occasional byproduct^18^. The results of this study imply that distance predictions contain information about residue pair flexibility and thus different protein conformers.

## Methods

### Dataset

The Database of protein Conformational Diversity in the Native State (CoDNaS) stores multiple PDB structures of single protein sequences.^34^ For our analysis the maximum RMSD pairs dataset of Monzon *et al.*^35^ was used. This CoDNaS subset contains 4791 pairs of PDB structures that had the maximum C_*α*_ RMSD amongst the pairwise comparisons between all conformers of a given protein. This subset includes only proteins that had at least five solved structures in the CoDNaS database to increase the reliability of the approximation of conformational diversity. Only X-ray structures with a resolution equal or less than 2.5Å were considered.

For our analysis a subset of the maximum RMSD pairs dataset was used which contained 3075 proteins. This subset contains only proteins that have no intrinsically disordered regions (unresolved patches of five or more residues) in any of the conformers. The list of PDB structure pairs can be found in SI table 1. PDB structures were pre-processed with clean pdb.py to remove alternative locations (downloaded from https://github.com/harryjubb/pdbtools/blob/master/clean_pdb.py on 14/11/2019).

For 2947 out of the 3075 proteins distance predictions, secondary structure assignment and chemical interaction analysis for both PDB structures could successfully be performed. These protein pairs constitute our analysis set and are marked with a 1 in the ‘analysis complete’ column of SI table 1.

### Co-evolutionary distance prediction

DMPfold^18^ was downloaded from https://github.com/psipred/DMPfold on 01/10/2019. In its normal application it generates distance predictions for all residue pairs and then feeds those into a structure modelling program (CNS). After model building the distance predictions are updated considering the built model (by default this step is done twice but can be varied). DMPfold was run with default parameters but without any model building and updates of distance predictions. Thus, distance predictions used in this work always refer to the initial distance prediction generated before model building iterations. Input features for the initial distance prediction were generated with hhblits 3.0.3 (multiple sequence alignment) against uniclust30 (2018 08).

Trivial contacts/residue pairs are defined as those between residues that are four or less residues in sequence apart. No distance prediction is generated by DMPfold for those residue pairs.

A distance prediction for a pair of residues refers to the predicted C_*β*_-C_*β*_ distance (in case of glycine C_*α*_) between those two residues and is termed predicted distance distribution here. Each predicted distance distribution is a vector of 20 points, representing the probabilities for each of the distance bins that DMPfold was trained to predict for a given sequence (or multiple sequence alignment). The first bin is ranging from 3.5-4.5Å, followed by seven bins of 0.5Å width up to a distance of 8Å and eleven bins of 1Å width up to 19Å. The last bin contained the probability density for distances over 19Å.

Target sequence of a distance prediction was the sequence derived from PDB structure 1 (see ‘pdb_id_1’ and ‘chain_id_1’ columns of SI table 1) through PDBParser(permissive=0) from Biopython 1.74^39^.

### Local maxima analysis

Distance probability distributions were analysed with the ‘peakdet’ function from Eli Billauer, version 3.4.05 (downloaded from https://gist.github.com/endolith/250860 on 18/10/2019). The function considers a point a local maximum if it has the local maximal value, and was followed (to the right) by a value lower by at least DELTA, which was chosen to be 0.03 (to maximise the fraction having two peaks while allowing at most three peaks. Detecting a local minimum (analogue definition with greater by at least delta) resets the local maximum. Thus, multiple local maxima can be found for each probability distribution. The last bin is never considered to be a maximum.

### Residue pair analysis

We only investigated non-trivial residue pairs that fulfilled the following true contact definition (true positives) and pairs that fulfilled the binary contact definition in at least one PDB structure if no local maximum could be detected at all (false negatives).

#### Definition of true contacts

The standard threshold for ‘binary’ contact predictors defines a residue pair to be in contact if the C_*β*_ C_*β*_ distance (in case of glycine C_*α*_) of two residues is 8Å or less^28^. Adapting this to distance predictions and multiple conformers, we define a true positive contact having at least one local maximum below 8Å and the ‘binary’ contact definition being satisfied in at least one of the conformers.

#### Flexibility definition

Residue pairs were classified into two distinct classes, rigid and flexible pairs; C_*β*_-C_*β*_ distances (C_*α*_ for glycine) and chemical bonds were used for this classification (Table 2).

**Table 2.** Flexibility class definitions

|  | Rigid residue pairs | Flexible residue pairs |
| --- | --- | --- |
| $C_\beta$-$C_\beta$ distance difference | < 1Å | >= 2Å |
| Chemical bond category | 1 + /1+ | 0/1+ |

#### C_β_-C_β_ distance difference

The absolute difference between the C_*β*_-C_*β*_ distances of the two structures of the same sequence was determined for each residue pair (in case of glycine C_*α*_). The absolute difference in distance had to be smaller than 1Å for the rigid classification and greater or equal to 2Å for the flexible classification.

#### Chemical bond analysis

To increase the confidence in a residue pair’s assignment to a flexibility class (rigid/flexible), we determined the presence of at least one chemical bond between those two residues in both PDB structures. A chemical bond is defined as any of the CREDO interactions^40^ detected by Arpeggio^41^; proximal interactions excluded. Arpeggio is a programme that calculates and visualises interatomic interactions from protein structures. It was downloaded from https://github.com/harryjubb/arpeggio. At least one chemical bond had to be present in both PDB structures (1+/1+) to classify for the rigid set and at least one bond in one of the structures but none in the other (0/1+) to classify for the flexible set.

#### Secondary structure pair types

Secondary structure assignment was determined using DSSP^42^ on PDB structure 1 (see ‘pdb_id_1’ and ‘chain_id_1’ columns of SI table 1). DSSP outputs an 8-letter code which we converted into two categories: secondary structure element (S) or loop/coil region (L). ‘H’, ‘G’, ‘I’ and ‘E’ were assigned to S and ‘B’, ‘T’, ‘S’ and’’ were assigned to L following the definition of Marks *et al.*.^24^ The secondary structure pair type is constituted by both residues’ assignments, leading to three types: ‘S-S’ (within or between secondary structures), ‘S-L’ (between secondary structure and loop) and ‘L-L’ (within or between loops).

### Set comparisons

For the comparison of fraction distributions the Mann-Whitney U test (two-sided) was applied as implemented in the Stats module of SciPy version 1.3.1.

### Rigid loop set

Daggert *et al.*^38^ defined a set of loops that showed very little movement in MD simulations (average below 0.5Å). Lists of the loop definitions and PDB structures used can be found in SI table 2. We followed the set definition of Marks *et al.* (which excluded one original loop because of an uncertainty in labelling).^24^ The number of local maxima was determined for each predicted distance distribution of residue pairs involving one of the defined loop residues.

