## Supplementary material for "Co-evolutionary Distance Prediction for Flexibility Prediction": SI Figures 1, 2, 3 and 4

### Supporting Information
#### Co-evolutionary Distance Prediction for Flexibility Prediction

##### Differences in number of local maxima between rigid and flexible sets are statistically significant

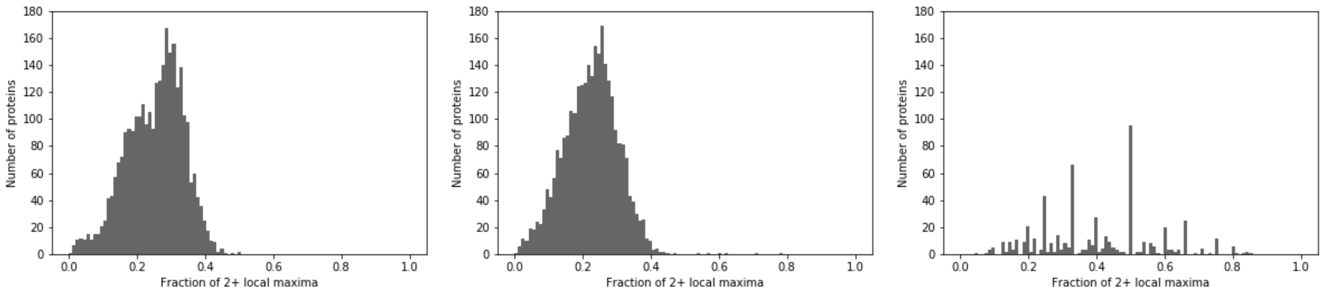


**(a)** All residue pairs **(b)** Rigid sets **(c)** Flexible sets

**Figure 1. Multiple local maxima occur significantly more often in flexible residue pairs than rigid residue pairs. (a)** Fractions of multiple local maxima for all residue pairs of a protein. **(b)** Fractions of multiple local maxima for the sets of rigid residue pairs. **(c)** Fractions of multiple local maxima for the sets of flexible residue pairs. As stated in the manuscript, all three distributions are significantly different from one another (Mann-Whitney U test): all vs. rigid sets with a p-value $< 10^{-44}$, all vs. flexible sets with a p-value $< 10^{-87}$ and rigid vs. flexible with a p-value $< 10^{-122}$.

#### Fractions of local maxima counts with a subset of unique CATH superfamily proteins

To check for a potential bias from overrepresentation of some CATH superfamilies in the dataset, we computed the fractions of local maxima counts for a subset of proteins with a maximum of one protein per CATH superfamily. If two or more proteins contained a (sub-)domain from the same CATH superfamily, only one of those proteins was selected at random.

For the sections of the main manuscript with plots of total, protein-averaged and secondary structure subgroup fractions of local maxima, plots were re-calculated with the unique CATH superfamily subset of the CoDNaS dataset. These plots and comparisons of this subset are qualitatively the same as the those of the full dataset.

##### Flexible residue pairs have more distance prediction local maxima than rigid residue pairs

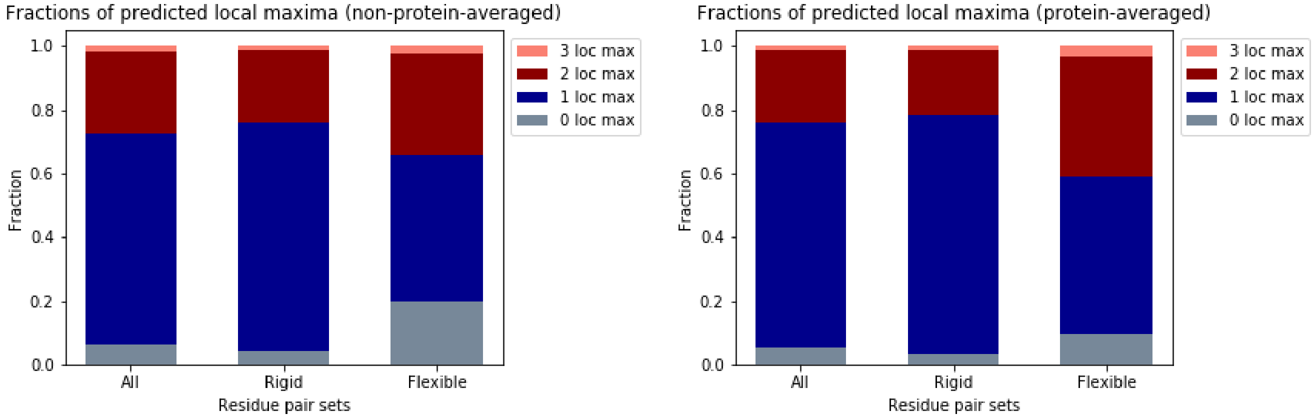


**(a)** Total fractions

**(b)** Protein-averaged fractions

**Figure 2. Fractions of local maxima counts are different between rigid and flexible residue pairs.** **(a)** Fractions of predicted local maxima across all classified residue pairs of all proteins, showing that flexible residue pairs are both more likely to have no local maximum or multiple local maxima. **(b)** Protein-averaged fractions of predicted local maxima, showing the same overrepresentation of no or multiple local maxima for flexible residue pairs.

##### Predicted distance distributions can capture flexibility independent of secondary structure

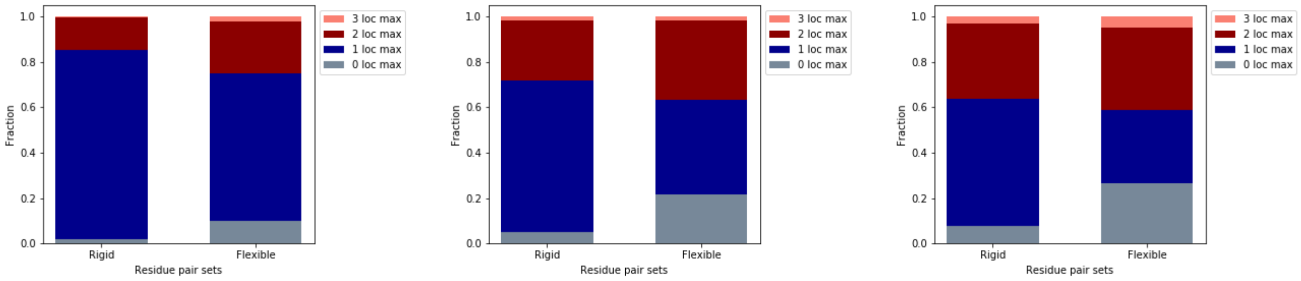


**(a)** S-S

**(b)** S-L

**(c)** L-L

**Figure 3. Predicted distance distributions capture flexibility independent of secondary structure.** Number of local maxima fractions were computed for subgroups depending on their secondary structure annotation: **(a)** 'S-S' when both residues were found to be part of a secondary structure element, **(b)** 'S-L' when one of the residues was in a secondary structure element and the other in a loop, and **(c)** 'L-L' when both were part of a loop/coil region. The different fractions of multiple local maxima (red) between rigid and flexible residue pairs are not only driven by an imbalance of secondary structure element and loop residues: for example the S-S subgroup of the flexible set has a fraction of multiple local maxima twice as big as the rigid set's.

##### Non-machine learning contact predictors rank flexible residue pairs lower than rigid residue pairs

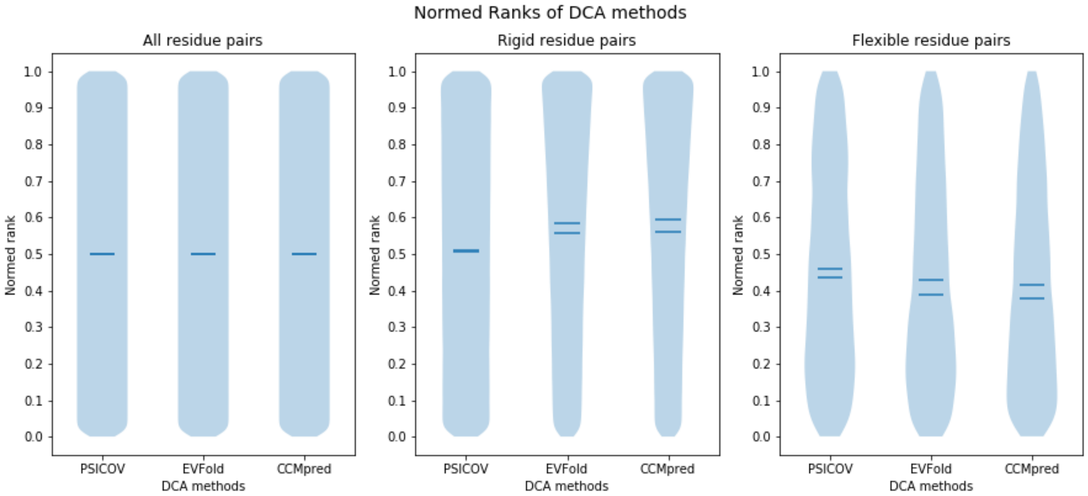


**Figure 4. Non-machine learning contact predictors based on direct coupling analysis (DCA) rank flexible residue pairs lower than rigid residue pairs.** All residue pairs is the set of residue pairs that are true contacts according to our definition for two PDB structures (see Methods). As the ranks in all three plots are normalised to just the true contacts of a given protein, ranks are uniformly distributed for All residue pairs and all three DCA methods. A rank of 1 indicates a prediction with the highest probability amongst all true contacts and a rank of 0 a prediction with the lowest probability. For rigid residue pairs predictions from EVfold and CCMpred are skewed towards 1 and predictions from PSICOV are close to uniformly distributed. For flexible residue pairs predictions from all three DCA methods are skewed towards 0 and means and medians are all below the ones for rigid residue pair predictions. This implies that flexible residue pairs are generally ranked lower (and thus predicted to be less likely in contact) than rigid residue pairs by all three investigated DCA methods.
